# Adipocytes in tumour microenvironment promote chemoresistance in TNBC through oxysterols

**DOI:** 10.1101/2025.04.24.650225

**Authors:** Alex Websdale, Yusur Al-Hilali, Baek Kim, Bethany J Williams, Laura Wastall, Hanne Roberg-Larsen, Thomas A. Hughes, Giorgia Cioccoloni, James L. Thorne

## Abstract

**Objective:** Triple-negative breast cancer (TNBC) patients with excess adipose tissue experience poorer disease-free survival than those with a healthy body mass index. Adipocytes, abundant in mammary tissue, store and release cholesterol, which can be hydroxylated to form oxysterols. These cholesterol derivatives activate the liver X receptor (LXR) pathway. This study tested the hypothesis that adipocytes contribute to an imbalanced tumour-microenvironment by exposing cancer cells to elevated oxysterols, mimicking chemotherapy-exposure conditions and priming for chemoresistance.

**Methods:** Tumour tissue microarray from 148 TNBC patients was assessed using immunohistochemistry for expression of CH25H, CYP46A1, CYP27A1 and P-glycoprotein, and survival outcomes assessed. Gene expression was compared between tumours from patients and mouse models with high versus low adiposity. In vitro, cells of the tumour-microenvironment were evaluated for oxysterol content, secretion, expression of relevant enzymes, and their ability to induce P-glycoprotein expression and drug resistance in TNBC cells.

**Results:** In patients, expression of oxysterol-synthesizing enzymes in stroma correlated with P-glycoprotein expression in cancer epithelial cells and was associated with shorter disease-free survival. Adipocytes conditioned media contained significantly higher oxysterols levels than that conditioned by other cell types and induced P-glycoprotein expression and drug resistance in MDA.MB.468 cells. Obese mice had elevated levels of P-glycoprotein in tumours compared to lean counterparts.

**Conclusions:** Adipocytes secrete oxysterols that promote drug resistance in vitro and correlate with oxysterol:Pgp axis and survival in vivo.

**Significance:** This study reveals a mechanism by which adipose tissue contributes to drug resistance in ER-negative breast cancers, identifying the oxysterol-Pgp axis as potential therapeutic target.

## Introduction

Triple negative breast cancer (TNBC) is a subtype of breast cancer defined by the lack of expression of oestrogen, progesterone and HER2 receptors. The absence of these receptors, which when expressed in other subtypes can be targeted with systemic hormone therapy or anti-HER2 therapy, means that these cancers can only be systemically treated with cytotoxic chemotherapy. Drug resistance is significant challenge and disease relapse can occur rapidly(Dent et al., 2007). The mechanisms of drug resistance in TNBC are complex and given disease heterogeneity, remain poorly understood. Overweight TNBC patients treated with chemotherapy have shorter disease-free survival (DFS) and overall survival (OS) compared to lean patients (Harborg et al., 2021) suggesting an interaction between adipose tissue and systemic chemotherapy.

The extensive heterogeneity of the TNBC microenvironment (TME) and the tumour’s mutational profiles make predicting which patients will suffer with drug resistance challenging. A high stroma to tumour ratio is associated with worse DFS particularly for TNBC patients (Millar et al., 2020, Kramer et al., 2019). The TME typically consists of immune cells, and stromal cells such as fibroblasts, which mount a cytoprotective interferon-b response to chemotherapy exposure (Millar et al., 2020, Broad et al., 2021). Adipocytes also comprise a significant proportion of the breast cancer TME. Adipose tissue contains up to 25% of the body’s total cholesterol rising to as much as 50% in obese individuals (Krause and Hartman, 1984). For this reason, overweight and obese patients breast tumours are particularly highly exposed to adipocytes cholesterol stores (Krause and Hartman, 1984). Circulating cholesterol is typically found in dynamic equilibrium with adipose cholesterol stores and elevated cholesterol levels at diagnosis is associated with lower disease-free survival (Rodrigues dos Santos et al., 2014)) in a manner that cholesterol metabolism should be considered as a therapeutic target alongside with chemotherapy.

Oxysterols are endogenous liver x receptor alpha (LXRa) ligands (Janowski et al., 1999, Janowski et al., 1996), which are derived from cholesterol by a single enzymatic hydroxylation reaction on cholesterol’s side-chain; their further metabolism leads to production of bile acids, seco-steroids, steroid hormones, and other important endocrine and paracrine factors (Slenter et al., 2018). Oxysterols are found in in breast cancer secreted exosomes (Roberg-Larsen et al., 2017) and in both ER-negative and ER-positive primary breast tumours at concentrations that activate LXR signalling (Solheim et al., 2019). ER-negative breast cancers are significantly more responsive to oxysterol signalling compared to other subtypes (Hutchinson et al., 2019) and in TNBC, expression of LXRα splice variants that lack the oxysterol binding domain is associated with longer DFS in TNBC (Lianto et al., 2021). Oxysterol driven LXRα activity induces expression and function of the P-glycoprotein (Pgp) efflux pump and is associated with chemoresistance in TNBC patients (Hutchinson et al., 2021). Here we have tested the hypothesis that the proximity and integration of adipose tissue, means that the TNBC microenvironment is chronically exposed to elevated levels of oxysterols which confer chemotherapy resistance in triple negative breast cancer via Pgp.

## Methods

### Cell lines

All cell lines were routinely maintained at 37 °C with 5% CO2 and tested for mycoplasma every 6 months. TNBC cell lines MDA.MB.468 and MDA.MB.453 were obtained from ATCC and were routinely passaged in Dulbecco’s Modified Eagle Medium (DMEM) Glutamax (DMEM, Thermo Fisher, Cat: 31966047), and supplemented with 10% FCS (Thermo Fisher, UK, Cat: 11560636) or when indicated for oxysterol pellet analysis, were instead cultured in CHUB-S7 media (see below). Generation (Hutchinson and Thorne, 2019) and application (Hutchinson et al., 2019) of MDA.MB.468 LXRα luciferase reporter cells has been described previously. Human pre-adipocyte CHUB-S7 stem cells, donated by Nestlé Research Centre, were originally isolated from subcutaneous abdominal adipose tissue of a 33-year-old obese female patient and immortalised by over-expressing hTERT as previously described (Darimont et al., 2003). CHUB-S7 were maintained in DMEM/Nutrient F12 Hams, supplemented with 10% FCS and 2 mM glutamine (Sigma, Cat: G7513). CHUB-S7s were differentiated into mature adipocytes (mAdipo) as previously described (Darimont et al., 2006) using differentiation medium supplemented with rosiglitazone (Cayman Chemical, Cat: 71740) and 100 U/ml penicillin/streptomycin (Sigma, Cat: P0781).

### Drugs and reagents

Epirubicin (Cayman, Michigan, US, Cat: 12091) was diluted in nuclease free water and protected from light. 24S-hydroxycholesterol (24OHC), 25-hydroxycholesterol (2OHC), and 27-hydroxycholesterol (27OHC, also known as 26-hydroxycholesterol and 25R26-hydroxycholesterol), together with internal standards (25-hydroxycholesterol-d6 and 27-hydroxycholesterol-d6) were from Avanti Polar Lipids (Alabaster, AL, USA). Stock solutions and calibration solutions were prepared as described in (Solheim et al., 2019). TaqMan assays (Thermo Fisher, Paisley, UK, Cat: 4331182): Pgp/*ABCB1* [Hs00184500_m1], *ABCA1* [Hs01059137_m1], *CYP27A1* [Hs0016803_m1], *CYP46A1* [Hs1042347_m1], *CH25H* [Hs04187516_m1] and *HPRT1* [Hs02800695_m1]. All stocks were stored at −20 °C. Antibodies used for IHC were: Pgp (Santa Cruz Biotech, California, US, Cat: sc73354), CYP27A1 (Abcam, Cambridge, UK - Cat: ab126785), CYP46A1 (Abcam, Cambridge, UK - Cat: ab198889), CH25H (Bioss, MA, US - Cat: bs6480R) and were validated previously or as previously described (Hutchinson et al., 2021).

### Conditioning of culture media by adipocytes and cancer associated fibroblasts

Mature adipocyte conditioned media (mAdipo-CM) was generated by adding 15mL of DMEM/F12 10% Charcoal stripped FBS (CS-FBS) into a confluent T75 and incubating for 48h. MDA.MB.468 CM (vehicle) was generated by seeding 4.5×10^6^ cells/mL into a T75 adding 15mL of DMEM/F12 containing CS-FBS, incubating for 48h. CM was centrifuged upon collection for 5 minutes at 200 rcf and used fresh or stored at -80°C until use. Then, MDA.MB.468 cells were seeded in 6 well plates at 6×10^5^cell/well for 4h and exposed to vehicle or mAdipo-CM at different ratios and top up with fresh media to reach the volume of 2mL/well for 16h, 24h and 48h. At the end of each treatment, cells were trypsinized, washed in PBS, and then suspended in different buffers to be assessed for luciferase expression, MTT or mRNA expression.

CAF-CM was generated by adding 10 mL of DMEM containing 10% FBS into a confluent T75 and incubating for 44h. To match the pH of the CAF-CM, MDA.MB.468 CM was generated by seeding 4.5×10^5^ cells/mL and conditioning DMEM containing 10% FBS for 24 h. All media was centrifuged upon collection for 5 minutes at 200rcf and stored at -80°C until use.

### Cell viability assay

For cell proliferation assay, MDA.MB.468 cells were treated with vehicle or mAdipo-CM as previously described. Cells were then trypsinized, washed and suspended in 200µL of PBS, and distributed in to a 96 well plate (90µL/well). For cell drug resistance assay, 2×10^4^/well MDA.MB.468 cells were seeded in a 96 well plate for 4h and then treated with either vehicle CM, or mAdipo-CM to reach the volume of 200µL/well for 24 or 48h respectively. For chemoresistance assay, epirubicin (Cayman Chemical, Michigan, US, Cat: 12091) was then added at different concentrations for 48h. Cells were washed and MTT reagent was added at final concentration of 0.5 mg/mL. In all experiments, after 4 h incubation at 37 °C, MTT solution was removed and replaced with 100μL of DMSO/well. Absorbance at 540nm was read using a CLARIOstar Plus microplate reader.

### Luciferase assay

Luciferase assays were performed pelleting 100 µL of MDA.MB.468 cells and suspending them in 100uL of Reporter Lysis buffer (Promega, UK, Cat: E1500). 20µL of cell lysate was distribute in to white-walled 96-well plates and luciferase expression measured using the Luciferase Assay system (Promega, UK, Cat: E1500) through luminescent signal using CLARIOstar Plus microplate reader (BMG LABTECH, Germany). Results were subsequently normalized using cell proliferation data to eliminate the proliferative effect of mAdipCM on TNBC cells, which could have distorted the luciferase data.

### Analysis of gene expression

Analysis of gene expression was performed as described previously (Hutchinson et al., 2019). Briefly, Reliaprep Minipreps for cell cultures (Promega, UK, Cat: Z6012) was used to extract mRNA, and GoScriptTM (Promega, UK, Cat:A5003) was used for the cDNA synthesis. Taqman Fast Advanced Mastermix (Thermo Fisher, Paisley, UK, Cat: 4444557) was mixed with Taqman assays and analysed in QuantStudio Flex 7 (Applied Biosystems Life Tech, Thermo Scientific,UK).

### Immunohistochemistry and adipocyte scoring

IHC was performed as previously described (Hutchinson et al., 2021). Tissue microarray (TMA) blocks sections were dewaxed in xylene and dehydrated in ethanol. For Pgp, CYP46A1 and CH25H antibodies, antigen retrieval was not performed. For CYP27A1 antibody, slides were heated in citrate buffer. Endogenous peroxidase activity was blocked with 0.3% H_2_O_2_ in methanol, and for Pgp, CYP46A1 and CH25H, a further block step was carried out using Blocker™ Casein in TBS (37532, Thermo Fisher Scientific, UK). Slides were incubated with antibodies for 1 h. Staining was visualised using secondary antibodies and SignalStain DAB substrate kit (Cell Signalling Technology, Cat: 80595). Nuclei were stained with Mayer’s haematoxylin and washed with Scott’s tap water. Sections were dehydrated with ethanol, washed with xylene and mounted onto coverslips with DePeX (Fluka Immunohistochemical staining within stromal regions was quantified using positive pixel count version 9 algorithm (Aperio Technologies, US) for ImageScope. Briefly, positive staining of Pgp, CYP27A1, CH25H and CYP46A1 was assessed semi-quantitatively using a weighted histoscore system. Immunohistochemical staining within stromal regions was quantified using positive pixel count version 9 algorithm (Aperio Technologies, US) for ImageScope. Briefly, positive staining of Pgp, CYP27A1, CH25H and CYP46A1 was assessed semi-quantitatively using a weighted histoscore system. Each tumour was scored in triplicate, with three cores embedded per tumour. Histoscores between 0-300 were determined, with this value calculated with the formula (1 x % of tumour cells weakly stained) + (2 x % of tumour cells moderately stained) + (3 x % of tumour cells strongly stained). Weighted histoscores were calculated for total stroma of the three enzymes and Pgp. Tumours were ignored if two or more duplicate cores were missing. For scoring of TMA sections, whole cores were scored. Tumours were ignored if two or more duplicate cores were missing. Accuracy of primary observer histoscores was previously reported (Hutchinson et al., 2021).

Stromal proportions of tumour cores were measured using ImageScope version 12.4.3 (Aperio Technologies, US) by drawing around the perimeter of the core, excluding necrotic regions and drawing around cancer regions in the core. Cancer area was then subtracted from the total area to find the stromal area of the core. Stromal area was then divided by total area to find the stromal proportion of the tumour.

Adipocyte count was performed on each tumour and scored in triplicate, with three cores embedded per tumour. Tumours were ignored if two or more replicate cores were missing. Accuracy of primary observer were verified by an independent observer for all tumour cores. Intraclass correlation was based on McGraw and Wong (McGraw and Wong, 1996) and Shrout and Fleiss convention (Shrout and Fleiss, 1979) and calculated using two-way mixed effects, consistency single rater method. Presence of adipocytes in each tumour was classified as 0 if none of the cores for each tumour was presenting adipocytes, or ≥1 if at least one core for each tumour presented adipocytes.

### Human samples: ethical approval, collection, and processing

Ethical approvals were obtained from Leeds (East) REC (reference numbers: 06/Q1206/180, 09/H1306/108). Patient cohort and clinical-pathological features have been described previously (Hutchinson et al., 2021) and updated for adipocytes count (**ST1**). Briefly, selection criteria: patients with TNBC/basal like tumours who had not undergone neoadjuvant therapy, tumour with sufficient stroma, absence of heavily necrotic or inflammatory regions, and whether resection blocks were available for normal tissue. Haematoxylin/eosin stain was used to identify suitable area for inclusion in the tissue microarray (TMA). Tumour cores of 0.6mm were taken in triplicate from 148 tumour samples and transferred into recipient wax blocks.

### Oxysterol quantification

MDA.MB.468, MDA.MB.453, CHUB-S7 pre-adipocyte, mature adipocyte and CAF pellets (3×10^5^ cells) were prepared as previously described (Roberg-Larsen et al., 2014) with some modifications. Cell pellets were added 100µL of 1.5 nM internal standard solution of deuterated 24OHC, 25OHC and 26OHC (Avanti Polar Lipids, Alabaster, AL, USA) in 2-propanol and shaken well before evaporation to dryness using a speedvac. The pellets were then re-dissolved in 20 µL 2-propanol before derivatization with Girard T reagent. Cell culture media samples (200μL) were prepared as previously described (Kømurcu et al., 2023) with some modifications: aliquots of 100 or 200µL were mixed with 100µL of 1.5nM internal standard solution of deuterated 24OHC, 25OHC and 26OHC (Avanti Polar Lipids, Alabaster, AL, USA) in 2-propanol and shaken well before evaporation to dryness using a speedvac. The residues were re-dissolved in 20µL isopropanol before derivatization with Girard T reagent. All samples had a final internal standard concentration of 200pM and were stored at 4 °C before analysis.

All cell and medium samples were analysed within a week using Dionex Ultimate 3000 UHPLC coupled to a TSQ Vantage triple quadrupole mass spectrometer (Thermo Scientific) as previously described (Solheim et al., 2019). Removal of excess derivatization reagents and other particular matter (sample clean-up) was performed on-line using an automatic filtration and filter back-flush (AFFL) solid phase extraction (SPE) system.

### Analysis of publicly available datasets

Publicly deposited data were obtained to compare TNBC tumours between obese and lean mice (Bousquenaud, 2022) and patients (Toro et al., 2016); details of experimental procedures are available in the corresponding citations. From the mouse dataset (GSE151866) we extracted gene expression data for lean (n=4) and obese (n=4) mice. For human samples (GSE78958) we extracted gene expression data and clinical information to select patients with basal-like breast cancer (n=98) and then classify by body mass index (BMI) as normal-weight (BMI<25: n=30), overweight (BMI 25-29.99: n=34), or obese (BMI≥30: n=34).

### Statistics

Analysis of protein correlations were assessed using Pearson’s rank. Kaplan-Meier analysis with 95% confidence intervals was used for survival curves, and log-rank test was performed to assess significance. One-way ANOVA with multiple comparisons was used to compare oxysterol content, RNA expression of oxysterol synthesising enzymes between mAdipo, CAF and breast cancer cell pellets, and between mAdipo-CM,CAF-CM and breast cancer cells CM. One-way ANOVA with multiple comparisons testing was also used to compare LXR transactivation and gene expression. TMA samples containing adipocytes were compared with samples without adipocytes using unpaired parametric one-tailed t-test. Spearman correlation was used to observe associations between gene expression the different BMI groups from GSE78958. One-tailed unpaired t-test was used to compare gene expression between lean and obese mice from GSE151866 and MDA.MB.468 gene expression following pre-treatment with mAdipo-CM. One-tailed paired t-test was used to analyse MTT assay of epirubicin dose response following pre-treatment of MDA.MB.468 with mAdipo-CM.

## Results

### Protein level of oxysterol synthesising enzymes in non-cancer compartment of the tumour microenvironment is associated with Pgp protein expression in cancer cells and with disease-free survival

To determine whether expression of stromal oxysterol enzymes was associated with Pgp expression in primary TNBC tumours and disease-free survival (DFS) of patients we analysed 148 TNBC tumour cores using a tissue microarray (TMA), details of which have been reported previously (Hutchinson et al., 2021). The proportion of the tumour found to be stromal correlated weakly, but significantly, with epithelial expression of Pgp protein (R^2^ = 0.09; p=0.0007). When patients were dichotomised into having high or low stroma-epithelial ratios using receiver operating characteristic (ROC curves shown in **SF1**), a high stroma to cancer cell ratio was, as expected, associated with shorter DFS (p=0.017) (**Fig1a**). Next, protein levels of the enzymes that convert cholesterol into 24OHC (CYP46A1), 25OHC (CH25H), or 27OHC (CYP27A1) was measured in the non-cancer cell fraction of the tumour (**Fig1b-d**). Non-cancer cell expression of all three enzymes was positively correlated with cancer cell expression of Pgp (CYP46A1: R^2^=0.22, p<0.0001; CH25H: R^2^=0.35, p<0.0001; CYP27A1: R^2^=0.07, p=0.0072). Elevated protein expression was associated with significantly shorter DFS for CYP46A1 (p=0.0075), CH25H (p=0.0024) and CYP27A1 (p=0.0183). These correlations and survival curves are consistent with the hypothesis that chemoresistance in TNBC cancer cells could at least in part be driven by oxysterols synthesised and secreted by the tumour microenvironment.

**Figure 1.**
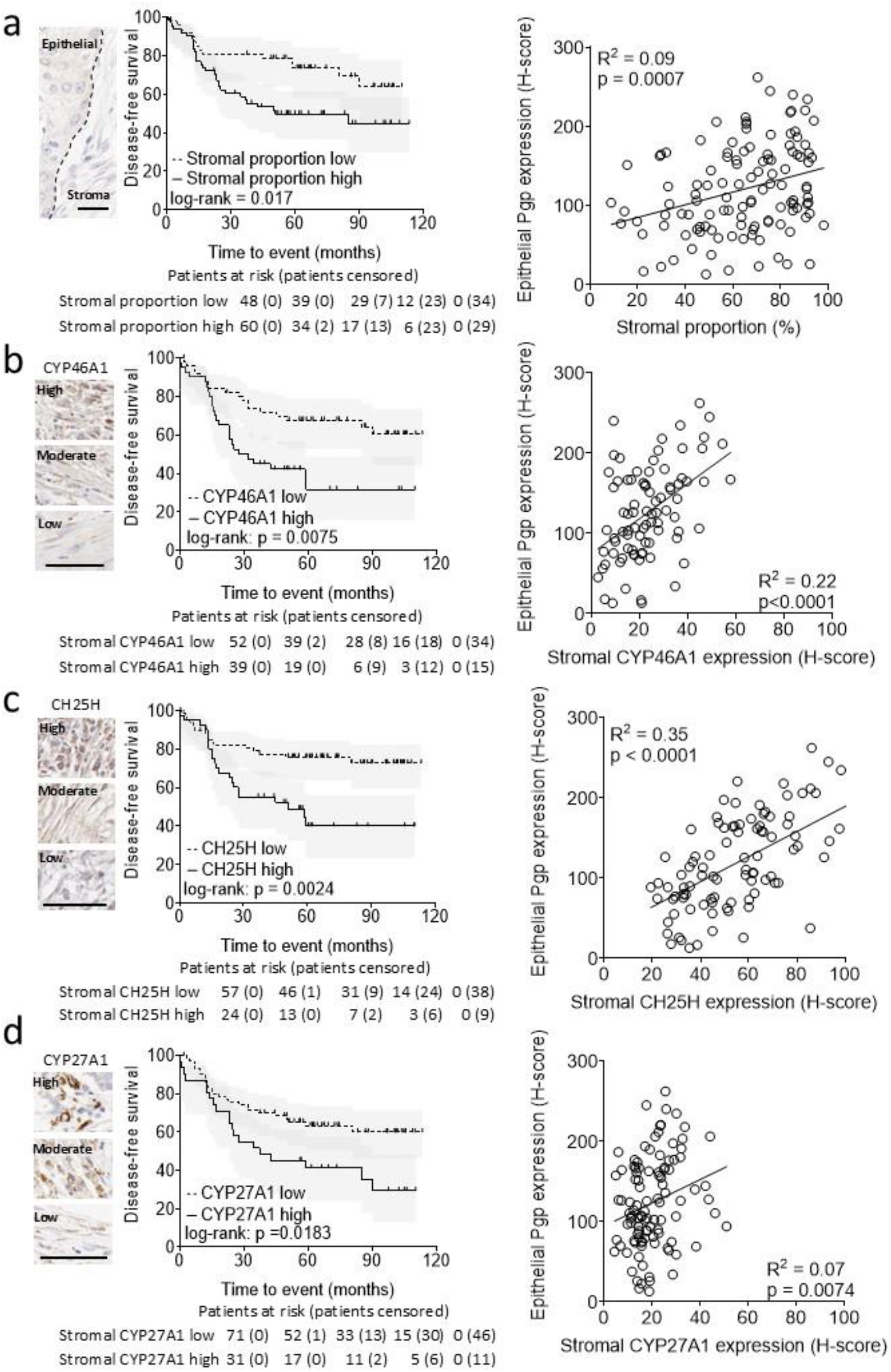
Expression of oxysterol synthesis enzymes in the triple negative breast cancer microenvironment is associated with elevated P-gp expression and shorter disease-free survival. From left to right, shown are representative images of tumour sections with 25mm scale bars (left), Kaplan-Meier survival curves (middle), and correlation analysis with cancer epithelial Pgp expression (right). Analyses shown are for epithelial to stromal proportions (a), CYP46A1 (b), CH25H (c) and CYP27A1 (d). Kaplan-Meier is shown with 95% confidence intervals with log-rank test with patients at risk of suffering an event are shown beneath. Spearman’s rank was used to assess correlation.

### Oxysterol synthesis and export by cells of the tumour microenvironment is dominated by adipocytes

The TME comprises a mixture of tumour cells and non-tumour ‘host’ cells. To establish if the tumour’s oxysterol pool, and therefore LXR activity within the cancer cell compartment, could be shaped by the non-cancer compartment of the TME, we compared the ability of TNBC cell lines, cancer associated fibroblasts (Verghese et al., 2013), and adipocytes to synthesise and secrete oxysterols, and their ability to activate LXR in cancer cells. First, we measured mRNA expression of the enzymes that synthesise the OHCs previously reported to be present in primary breast tumour samples (Solheim et al., 2019). Synthesis of CYP46A1, the 24OHC synthesising enzyme, was significantly higher in mAdipo cells than all other cell types (**Fig2a**; p<0.0001 for all). For CH25H, which generates 25OHC, expression was also higher in mAdipo than CAFs, MDA.MB.453 and MDA.MB.468 cell lines (**Fig2b**; p=0.0568, p=0.0863, p=0.0721, respectively). Expression of CYP27A1 was similar between mAdipo and CAFs (**Fig2c**; p>0.05) but both had significantly more mRNA than the TNBC cell lines (p<0.001 for all). Regarding intracellular OHC concentrations, mAdipo had significantly higher levels than the TNBC cell lines for all three OHCs (**Fig2d-f**; p<0.05 for all), while CAFs had significantly less 24OHC than mApido (p=0.0035) and were statistically equivocal to the TNBC cell lines for 25OHC and 27OHC. To establish if oxysterols were secreted into the TME by the four cell types we sampled cell culture media after conditioning for 48hr types. 27OHC was the most abundant oxysterol released into the media by an order of magnitude in mAdipo and CAFs (**Fig2g**). mAdipo conditioned media contained significantly more OHC compared to CAFs, for 24OHC (7.8-fold, p<0.0001), 25OHC (11-fold, p=0.0134) and 27OHC (5.3-fold, p<0.0001) (**Fig2g**). CAF conditioned media had significantly less OHCs than mAdipo, and contained either the same or less OHCs than MDA.MB.453 and MDA.MB.468 conditioned media. Finally, to establish if the conditioned media contained sufficient oxysterols to activate LXR signalling in TNBC epithelial cells, we culture LXR reporter MDA.MB.468 cells in conditioned media. A modest but significant 2.4-fold increase in LXR reporter gene activity was observed after 24 hr culture with 100% mAdipoCM (p<0.0001). At 48 hr LXR reporter gene activity was induced in a manner dependent on dose of conditioned media (75% mAdipoCM: 2.8-fold; 100% mAdipoCM: 3.9-fold; p = 0.0018) (**Fig2h**). From these data we concluded that, of the TME components we could utilise in vitro, mAdipo produce significantly higher levels of oxysterols than other cell types.

**Figure 2.**
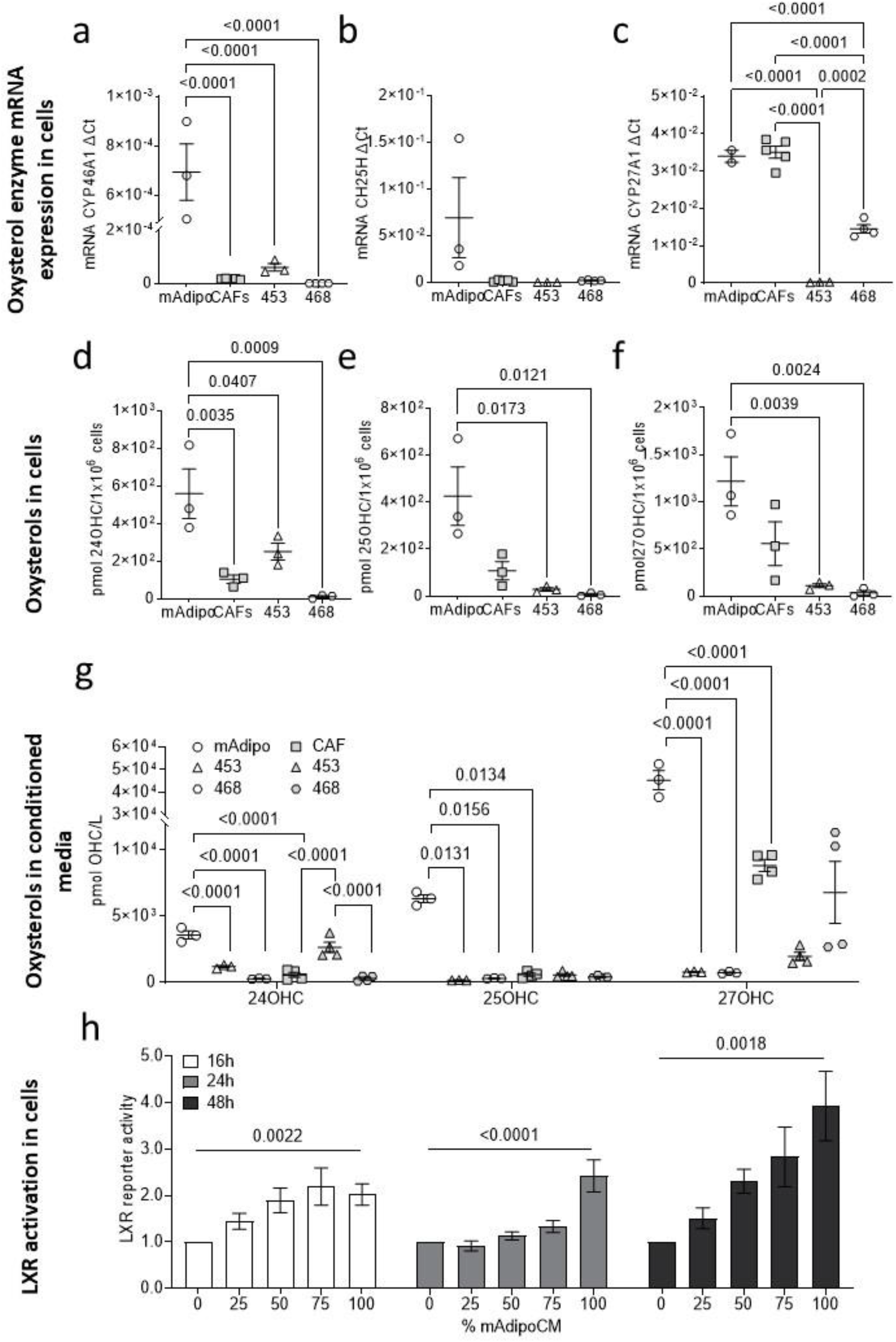
Mature adipocytes but not cancer associated fibroblasts contain and secrete higher concentrations of oxysterols relative to cancer epithelial cells. mRNA expression of CYP46A1 (a), CH25H (b) and CYP27A1 (c) from mAdip, CAFs, MDA.MB.468 and MDA.MB.453 cells. Oxysterol concentrations in pellets of 1×10^6 cells (d-f). Data shown are mean of 2-4 independent replicates with standard error and are analysed with one-way ANOVA. Media concentrations of 24OHC, 25OHC, and 27OHC (g) after conditioning with mAdip, CAFs, MDA.MB.468 and MDA.MB.453 cells. Cancer cell data are shown twice to match the different growth culture media required for mAdipo cells (charcoal stripped DMEM/F12) and CAFs (10% FBS DMEM). Data shown are mean of 3-4 independent replicates with standard error and are analysed with one-way ANOVA. Conditioned media from mAdipo drives LXR activity in MDA.MB.468 reporter cells (h). Data shown are mean of 7-9 independent replicates and are analysed with one-way ANOVA.

### Adiposity is associated with higher Pgp and CH25H expression in primary tumours

We next hypothesised that if adipocytes could secrete oxysterols, then adipocyte numbers should be associated with enhanced expression of LXR target genes. As we had already measured expression of the LXR target gene ABCB1/Pgp in the TMA cohort (**Fig1a**), we reinterrogated these tumour cores, by scoring them for the number of adipocytes present. In tumour cases where there was at least one visible adipocyte, Pgp expression was significantly higher (p=0.0388) compared to cores where no adipocytes were visible (**Fig3a**). We also found that there were significantly higher levels of another LXR target gene, the 25OHC synthesising enzyme CH25H (p=0.039) in cores with at least one adipocyte (**SFig2a**). Expression of other OHC synthesising enzymes CYP46A1 (**SFig2b**) and CYP27A1 (**SFig2c**) in cancer cells were not significantly different between adipocyte containing cores and non-adipocyte containing cores, which was expected as neither are known LXR target genes and thus serve as an internal negative control. No significant association was found between TMA samples with adipocytes presence and survival outcome (p= 0.1303)(**SFig3a**).

**Figure 3.**
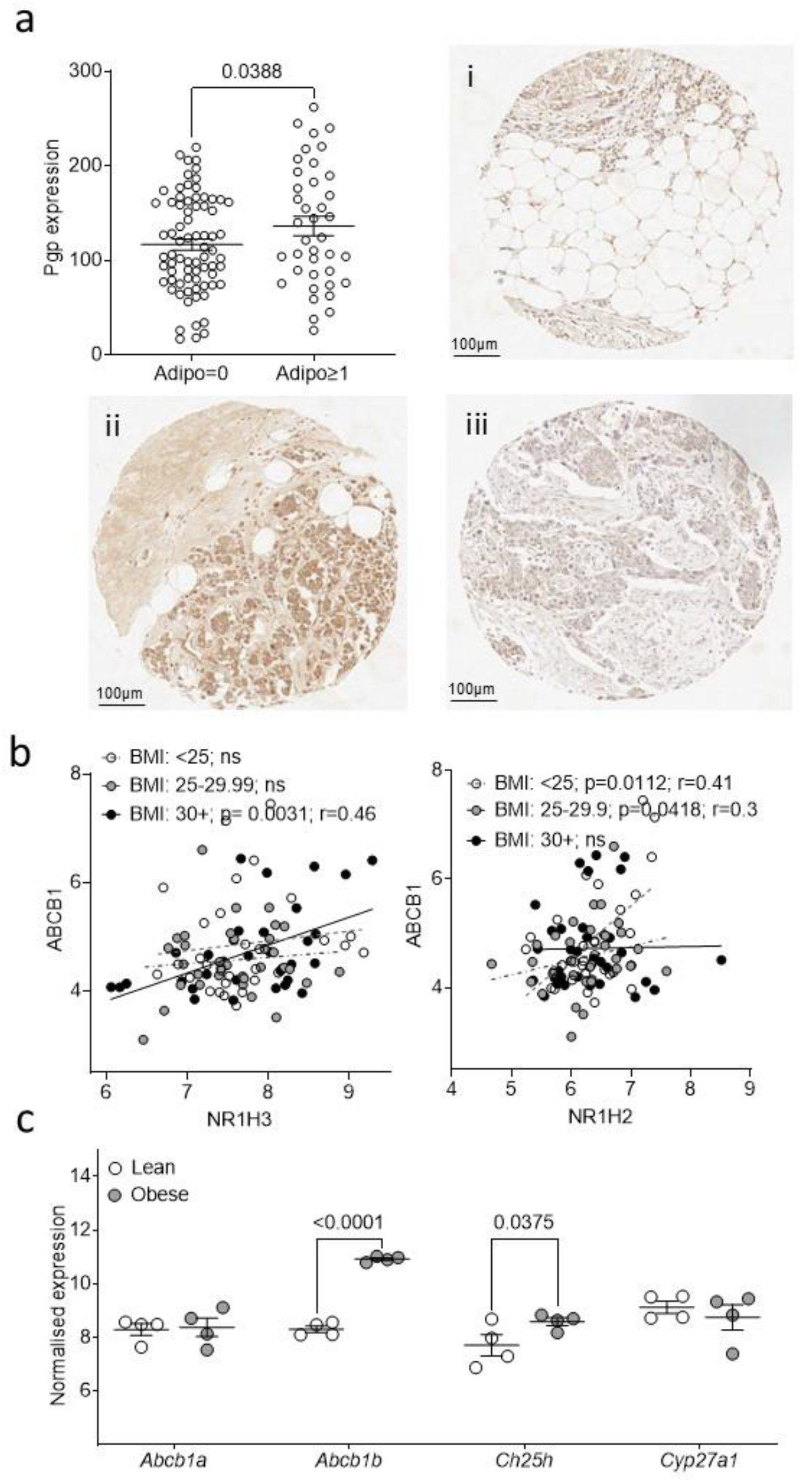
Obesity is associated with activity of the LXR:Pgp axis in TNBC patients and animal models. Presence of adipocytes is associated with elevated protein expression of Pgp (a) in a 148 TNBC patient tissue microarray. TMA samples are categorised as with or without adipocytes according to presence or absence of adipose cells in respective triplicates cores. Data shown are samples with (n=39) or without (n=74) adipocytes with available Pgp staining, and analysed using unpaired parametric one-tail t-test In obese but not overweight or normal weight TNBC patients (GSE78958), Pgp is significantly positively correlated with NR1H3 (b) Data shown are from patients with basal-like breast cancer (n=98) and grouped according to BMI information (<25 n=30; 25-29.99 n=34; ≥30 n=34) and analysed using Spearman correlation. Gene expression (GSE151866) of Pgp and CH25H is elevated in TNBC tumours from obese mice compared to lean mice (c). Data shown are from obese (n=4) and lean (n=4) with TNBC mice, and analysed using one-tailed unpaired t-test.

To validate this finding in additional datasets we performed a literature review and identified a publicly available dataset ((GSE78958 (Toro et al., 2016)) that contained basal-like cancers (n=98), which were annotated with categorical BMI data (<25, 25-29.99, 30+) and reported expression of LXR target gene ABCB1/Pgp. ABCB1 mRNA expression was positively correlated with NR1H3/LXRα (r=0.46, p=0.0031) but interestingly, only in obese patients (**Fig3b**). Notably, this correlation was not found in Luminal A or Luminal B subtypes from the same dataset (**SFig4**). Given the lack of Her2+ samples in the GSE78958 dataset the analysis on this subtype was not performed. In addition, we identified a publicly available transcriptomic dataset from diet induced obese mice implanted with the Py230 cell line model of basal-like breast cancer (GSE151866). In the Py230 tumours of the obese mice there was significantly higher levels of *Abcb1b* (p<0.0001) and *Ch25h* (p=0.036) compared to the tumours of lean mice (**Fig3c**).

### Adipocytes induce Pgp expression and confer chemoresistance in TNBC cells

Next, to determine if adipocytes could cause induction of Pgp in TNBC cells, mAdipo-CM was again harvested and added to MDA.MB.468 culture media. Culture of MDA.MB.468 cells with mAdipo-CM led to significant induction of ABCB1 gene expression (**Fig4a,b**; 24 hr: 1.5-fold, p=0.012; 48 hr: 1.3-fold, p=0.024) and enhanced survival in the presence of cytotoxic doses of epirubicin (**Fig4c,d**; p<0.05). These data are consistent with the hypothesis that an adipocyte secreted factor(s) drives expression of the Pgp drug efflux pump in cancer cells and confers chemoresistance.

**Figure 4.**
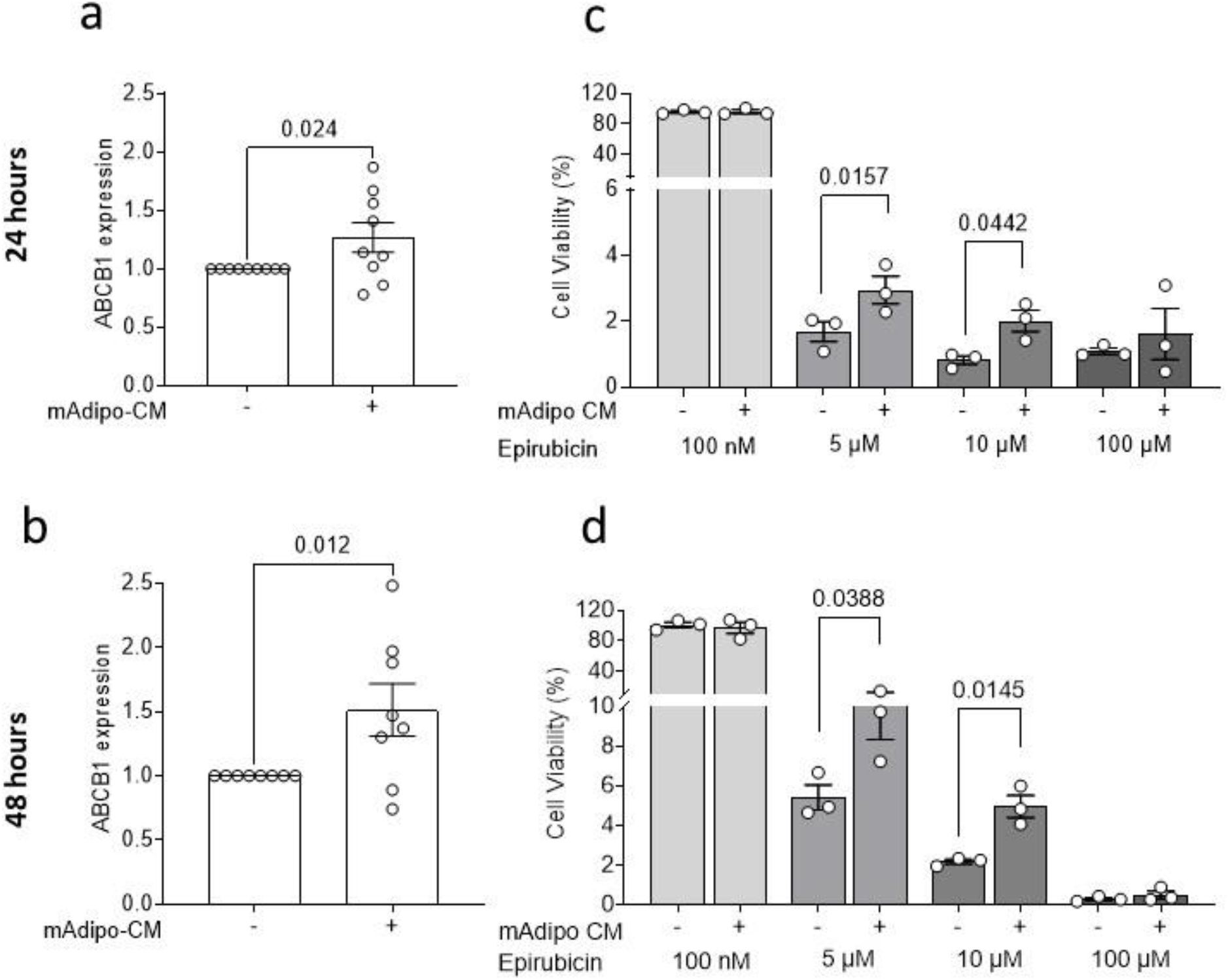
Adipocyte conditioned confers resistance to epirubicin. Induction of ABCA1 and ABCB1 after culture with mAdipo-CM for 24 hr (a) and 48 hr (b). Data show mean of 7-9 independent replicates with standard error. One-tailed t-tests were used to compare means. mAdipo-CM confers resistance to epirubicin after 24 hr (c) and 48 hr (d). Data are normalized to matched 0% conditioned media. Data show mean of three independent replicates with standard error. One-tailed t-tests were used to compare means.

## Discussion

The aim of this study was to test the hypothesis that the cholesterol store within adipocytes, that are proximal to breast tumour tissue, could provide a source of oxysterols that confer chemoresistance in TNBC. In vitro, adipocytes were found to contain and secrete relatively high levels of oxysterols and when TNBC cancer cells were cultured with adipocyte conditioned media they increased expression of the LXR target gene Pgp and had greater resistance to epirubicin. In patients, the expression of several oxysterol synthesising enzymes within the non-tumour component of the tumour microenvironment, were associated with elevated Pgp expression in the cancer cells and shorter disease-free survival for the patients. The amount of adipose cells present in the TME of our TNBC cohort was also significantly associated with expression of CH25H, which is not just responsible for synthesising the LXR ligand 25OHC, but is itself an LXRα target (Liu et al., 2018). CH25H gene expression was elevated in the tumours of obese mice relative to lean, as was Abcb1/Pgp. We are unable to determine if CH25H expression is correlated with Pgp because it is synthesising the necessary ligand for LXR driven transcription of the Pgp gene or because CH25H and Pgp are both under similar regulatory mechanisms; these are not mutually exclusive scenarios.

In previous work, ER-negative breast cancer was found to be more responsive to oxysterols and LXR signalling than ER-positive disease (Hutchinson et al., 2019), despite similar oxysterol concentrations between breast cancer subtypes (Solheim et al., 2019). Furthermore, activity of the LXR-Pgp axis in TNBC is linked to chemotherapy resistance and reduced patient survival (Hutchinson et al., 2021). If the OHC-LXR axis contributes to chemoresistance, attempts to reduce circulating cholesterol prior to commencing chemotherapy may dampen this resistance mechanism. However, this needs careful evaluation in the clinical trial setting. It is important to note that our data do not suggest that overweight/obesity *per se* exacerbate chemoresistance, but that conversion of cholesterol to oxysterols can occur independently of BMI. The proximity and abundance of adipose tissue in the breast cancer TME, coupled with TNBC being highly responsive to oxysterol signalling, may mean that sufficient oxysterols to contribute to chemoresistance are present in most triple negative breast tumours and thus represent an innate, yet potentially modifiable, form of resistance. Interventions that decouple the cholesterol-oxysterol pathway rather than just weight loss or cholesterol lowering, at least in the period between diagnosis and onset of neoadjuvant or adjuvant chemotherapy, may be most effective.

Our work has several important limitations. Firstly, the induction of Pgp gene expression observed after culture with conditioned media was low to moderate, which although is common for modulation of this efflux pump by nutritional or metabolic factors (Hutchinson et al., 2021) may not be sufficient to explain the chemoprotection we observed and other drug resistance pathways may have been induced. Although this induction was moderate, in the in vivo context, the local tumour cells are exposed for longer and to higher levels of oxysterols than we observed in conditioned media experiments. This could also mean that there are additional cell types contributing to the oxysterol pools in primary tumours that we have not recapitulated in our model. Although we found adipocytes to produce more oxysterols than cancer associated fibroblasts, this is in the context of a single immortalised (albeit it derived from a primary breast tumour) cell line and we did not attempt to confirm these findings in the primary setting. Our comparisons of enzyme expression therefore not be reproducible with other adipocyte or CAF sources, including analyses in primary material. These limitations should be addressed in future work because CAFs are able to confer chemoresistance in claudin-low TNBC (the work presented here is conducted exclusively in claudin-high TNBC cell lines) in an IFN-b dependent manner (Broad et al., 2021) and there is extensive cross talk reported between LXR and IFN pathways in several tissue types (Hao et al., 2009, Reboldi et al., 2014, Miao et al., 2016).

## Data Availability Statements

Data supporting the findings of this study are available from the corresponding authors upon reasonable request.

## Author Contributions

JLT sourced funding for salaries and consumables. JLT and GC wrote the manuscript. GC and AW performed the laboratory experiments. LW built and counter-scored the TMA for IHC. AW and BW scored the TMA IHC. GC and YAH performed the adipocyte counts. HRL and AW performed oxysterol analysis. JLT, TAH, and BK supervised the project. All authors approved the final manuscript.

## Acknowledgements

CHUB-S7 pre-adipocytes were kindly provided by Dr Christian Darimont from Nestlé Research Centre under Material Transfer agreement signed 18-10-2019.

## Graphical Abstract

**Adipocytes-derived oxysterols protect TNBC cells from chemotherapy activating LXRα-Pgp axis**. Figure created with Biorender.

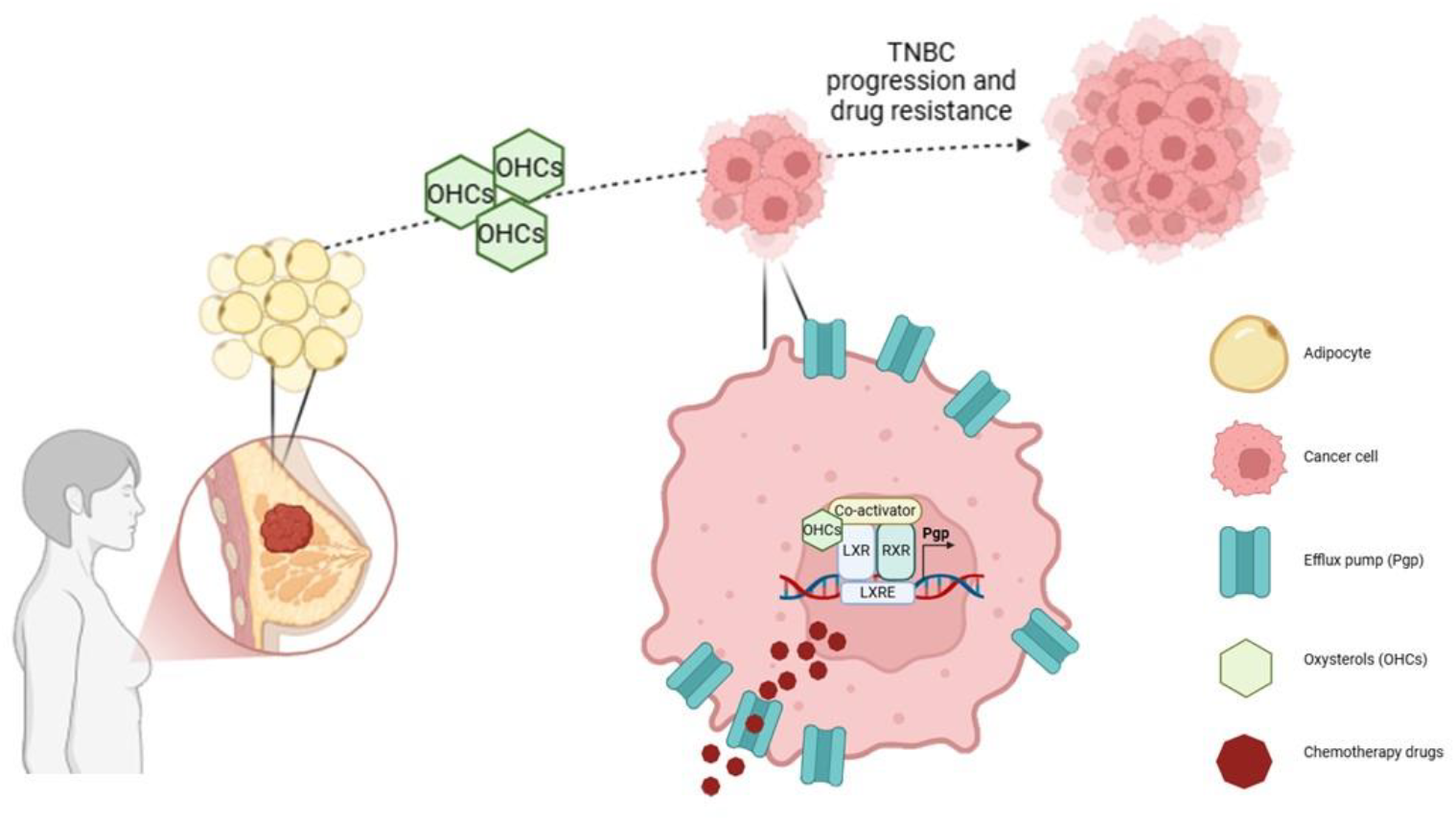

## References

Bousquenaud, M. D., I; RÜEgg, C. 2022. Tumor genes regulated in obesity-mediated breast cancer. In: BIOINFORMATICS, S. I. O. (ed.). NCBI, Gene Expression Omnibus.

Broad, R. V., Jones, S. J., Teske, M. C., Wastall, L. M., Hanby, A.M., Thorne, J. L. & Hughes, T. A. 2021. Inhibition of interferon-signalling halts cancer-associated fibroblast-dependent protection of breast cancer cells from chemotherapy. British journal of cancer, 124, 1110–1120.

Darimont, C., Avanti, O., Zbinden, I., Leone-Vautravers, P., Mansourian, R., Giusti, V. & Mace, K. 2006. Liver X receptor preferentially activates de novo lipogenesis in human preadipocytes. Biochimie, 88, 309–18.

Darimont, C., Zbinden, I., Avanti, O., Leone-Vautravers, P., Giusti, V., Burckhardt, P., Pfeifer, A. M. & Mace, K. 2003. Reconstitution of telomerase activity combined with HPV-E7 expression allow human preadipocytes to preserve their differentiation capacity after immortalization. Cell Death Differ, 10, 1025–31.

Dent, R., Trudeau, M., Pritchard, K. I., Hanna, W. M., Kahn, H. K., Sawka, C. A., Lickley, L. A., Rawlinson, E., Sun, P. & Narod, S. A. 2007. Triple-negative breast cancer: clinical features and patterns of recurrence. Clin Cancer Res, 13, 4429–34.

Hao, X. R., Cao, D. L., Hu, Y. W., Li, X. X., Liu, X. H., Xiao, J., Liao, D. F., Xiang, J. & Tang, C. K. 2009. IFN-gamma down-regulates ABCA1 expression by inhibiting LXRalpha in a JAK/STAT signaling pathway-dependent manner. Atherosclerosis, 203, 417–28.

Harborg, S., Zachariae, R., Olsen, J., Johannsen, M., Cronin-Fenton, D., Boggild, H. & Borgquist, S. 2021. Overweight and prognosis in triple-negative breast cancer patients: a systematic review and meta-analysis. NPJ Breast Cancer, 7, 119.

Hutchinson, S. A., Lianto, P., Roberg-Larsen, H., Battaglia, S., Hughes, T. A. & Thorne, J. L. 2019. ER-Negative Breast Cancer Is Highly Responsive to Cholesterol Metabolite Signalling. Nutrients, 11.

Hutchinson, S. A. & Thorne, J. L. 2019. A Stable Luciferase Reporter System to Characterize LXR Regulation by Oxysterols and Novel Ligands. Methods Mol Biol, 1951, 15–32.

Hutchinson, S. A., Websdale, A., Cioccoloni, G., Roberg-Larsen, H., Lianto, P., Kim, B., Rose, A., Soteriou, C., Pramanik, A., Wastall, L. M., Williams, B. J., Henn, M. A., Chen, J. J., Ma, L., Moore, J. B., Nelson, E., Hughes, T. A. & Thorne, J. L. 2021. Liver x receptor alpha drives chemoresistance in response to side-chain hydroxycholesterols in triple negative breast cancer. Oncogene, 40, 2872–2883.

Janowski, B. A., Grogan, M. J., Jones, S. A., Wisely, G. B., Kliewer, S. A., Corey, E. J. & Mangelsdorf, D. J. 1999. Structural requirements of ligands for the oxysterol liver X receptors LXRalpha and LXRbeta. Proc Natl Acad Sci U S A, 96, 266–71.

Janowski, B. A., Willy, P. J., Devi, T. R., Falck, J. R. & Mangelsdorf, D. J. 1996. An oxysterol signalling pathway mediated by the nuclear receptor LXR alpha. Nature, 383, 728–31.

KØMurcu, K. S., Wilhelmsen, I., Thorne, J. L., Krauss, S., Wilson, S. R., Aizenshtadt, A. & RØBerg-Larsen, H. 2023. Mass spectrometry reveals that oxysterols are secreted from non-alcoholic fatty liver disease induced organoids. The Journal of Steroid Biochemistry and Molecular Biology, 232, 106355.

Kramer, C. J. H., Vangangelt, K. M. H., Van Pelt, G. W., Dekker, T. J. A., Tollenaar, R. & Mesker, W. E. 2019. The prognostic value of tumour-stroma ratio in primary breast cancer with special attention to triple-negative tumours: a review. Breast Cancer Res Treat, 173, 55–64.

Krause, B. R. & Hartman, A. D. 1984. Adipose tissue and cholesterol metabolism. J Lipid Res, 25, 97–110.

Lianto, P., Hutchinson, S. A., Moore, J. B., Hughes, T. A. & Thorne, J. L. 2021. Characterization and prognostic value of LXR splice variants in triple-negative breast cancer. iScience, 24, 103212.

Liu, Y., Wei, Z., Ma, X., Yang, X., Chen, Y., Sun, L., Ma, C., Miao, Q. R., Hajjar, D. P., Han, J. & Duan, Y. 2018. 25-Hydroxycholesterol activates the expression of cholesterol 25-hydroxylase in an LXR-dependent mechanism. J Lipid Res, 59, 439–451.

Mcgraw, K. O. & Wong, S. P. 1996. Forming inferences about some intraclass correlation coefficients. Psychological methods, 1, 30.

Miao, C. M., He, K., Li, P. Z., Liu, Z. J., Zhu, X. W., Ou, Z. B., Ruan, X. Z., Gong, J. P. & Liu, C. A. 2016. LXRalpha represses LPS-induced inflammatory responses by competing with IRF3 for GRIP1 in Kupffer cells. Int Immunopharmacol, 35, 272–279.

Millar, E. K., Browne, L. H., Beretov, J., Lee, K., Lynch, J., Swarbrick, A. & Graham, P. H. 2020. Tumour Stroma Ratio Assessment Using Digital Image Analysis Predicts Survival in Triple Negative and Luminal Breast Cancer. Cancers (Basel), 12.

Reboldi, A., Dang, E. V., Mcdonald, J. G., Liang, G., Russell, D. W. & Cyster, J. G. 2014. Inflammation. 25-Hydroxycholesterol suppresses interleukin-1-driven inflammation downstream of type I interferon. Science, 345, 679–84.

Roberg-Larsen, H., Lund, K., Seterdal, K. E., Solheim, S., Vehus, T., Solberg, N., Krauss, S., Lundanes, E. & Wilson, S. R. 2017. Mass spectrometric detection of 27-hydroxycholesterol in breast cancer exosomes. J Steroid Biochem Mol Biol, 169, 22–28.

Roberg-Larsen, H., Lund, K., Vehus, T., Solberg, N., Vesterdal, C., Misaghian, D., Olsen, P. A., Krauss, S., Wilson, S. R. & Lundanes, E. 2014. Highly automated nano-LC/MS-based approach for thousand cell-scale quantification of side chain-hydroxylated oxysterols [S]. Journal of Lipid Research, 55, 1531–1536.

Rodrigues Dos Santos, C., Fonseca, I., Dias, S. & Mendes De Almeida, J.C. 2014. Plasma level of LDL-cholesterol at diagnosis is a predictor factor of breast tumor progression. BMC Cancer, 14, 132.

Shrout, P. E. & Fleiss, J. L. 1979. Intraclass correlations: uses in assessing rater reliability. Psychol Bull, 86, 420–8.

Slenter, D. N., Kutmon, M., Hanspers, K., Riutta, A., Windsor, J., Nunes, N., Melius, J., Cirillo, E., Coort, S. L., Digles, D., Ehrhart, F., Giesbertz, P., Kalafati, M., Martens, M., Miller, R., Nishida, K., Rieswijk, L., Waagmeester, A., Eijssen, L. M. T., Evelo, C. T., Pico, A. R. & Willighagen, E. L. 2018. WikiPathways: a multifaceted pathway database bridging metabolomics to other omics research. Nucleic Acids Res, 46, D661–D667.

Solheim, S., Hutchinson, S. A., Lundanes, E., Wilson, S. R., Thorne, J. L. & Roberg-Larsen, H. 2019. Fast liquid chromatography-mass spectrometry reveals side chain oxysterol heterogeneity in breast cancer tumour samples. J Steroid Biochem Mol Biol.

Toro, A. L., Costantino, N. S., Shriver, C. D., Ellsworth, D. L. & Ellsworth, R. E. 2016. Effect of obesity on molecular characteristics of invasive breast tumors: gene expression analysis in a large cohort of female patients. BMC Obes, 3, 22.

Verghese, E. T., Drury, R., Green, C. A., Holliday, D. L., Lu, X., Nash, C., Speirs, V., Thorne, J. L., Thygesen, H. H., Zougman, A., Hull, M. A., Hanby, A. M. & Hughes, T. A. 2013. MiR-26b is down-regulated in carcinoma-associated fibroblasts from ER-positive breast cancers leading to enhanced cell migration and invasion. J Pathol, 231, 388–99.

