## Supplementary Information for "Adipocytes in tumour microenvironment promote chemoresistance in TNBC through oxysterols"

**Supplementary table 1. Patient characteristics.**

| Characteristic | Category | Tissue Microarray<br>No. of patients = 148 (%) |
|---|---|---|
| **Tumour Grade** | 1 | 2 (1) |
| | 2 | 19 (13) |
| | 3 | 124 (84) |
| | N/A | 3 (2) |
| **Survival status** | Alive | 101 (68) |
| | Deceased | 47 (32) |
| **Recurrence status** | None | 102 (69) |
| | Present | 46 (31) |
| **Pgp** | | 125 (84) |
| **CYP46A1** | | 95 (64) |
| **CH25H** | | 97 (65) |
| **CYP27A1** | | 92 (62) |
| **Adipocytes** | Absent | 90 (61) |
| | Present | 52 (35) |
| | N/A | 6 (4) |

Clinicopathological features of the TMA cohort (n = 148).

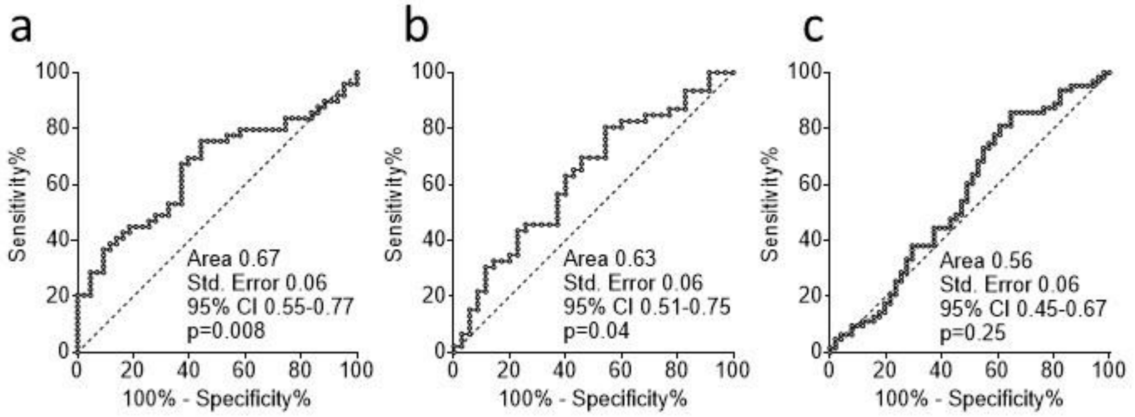


**Supplementary Figure 1: ROC curve used to determine cut offs used in Kaplan-Meier analysis.** Wilson-Brown method was used to calculate confidence intervals (95%) for sensitivity and specificity.

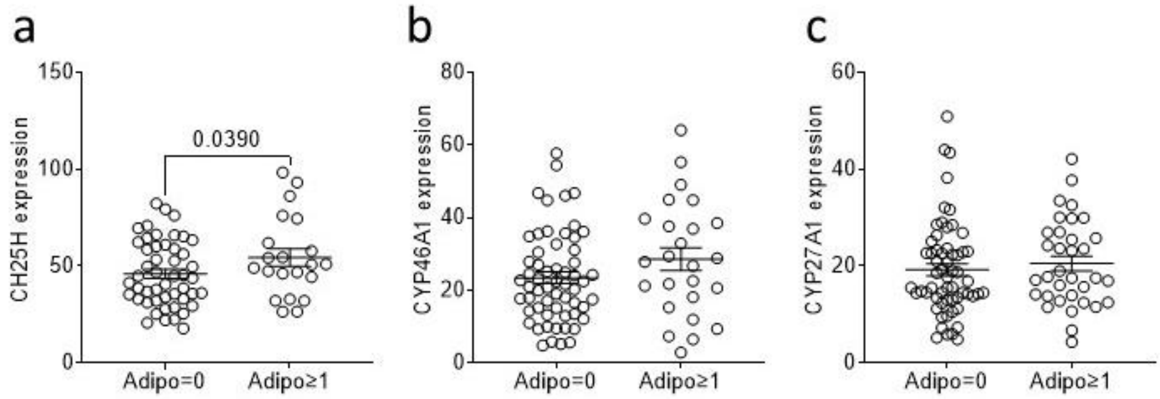


**Supplementary Figure 2: TME adipose cells are significantly associated with expression of CH25H in TNBC cohort.** Presence of adipocytes is associated with elevated protein expression of CH25H (a), but not CYP46A1 (b) or CYP27A1 (c) in a 148 TNBC patient tissue microarray. Data shown are samples with (n=39) or without (n=74) adipocytes with available CH25H staining and analysed using unpaired parametric one-tail t-test.

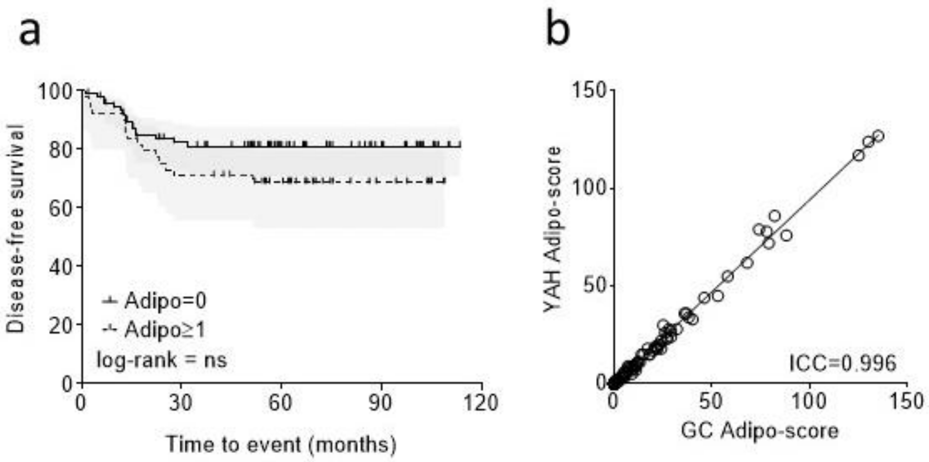


**Supplementary Figure 3: Adipocytes in TME do not associate with survival outcome.** Kaplan-Meier survival curves comparing survival related to presence/absence of adipocytes in a 148 TNBC patient tissue microarray (a). TMA samples are categorised as with or without adipocytes according to presence or absence of adipose cells in respective triplicates cores. Data shown are samples with (n=39) or without (n=74) adipocytes with available survival outcome data. Kaplan-Meier is shown with 95% confidence intervals with log-rank test with patients at risk of suffering an event are shown beneath. Intraclass correlation between cell counts from two independent scores for adipocytes (b). Scores given by scorer 1, GC, are plotted against scorer 2, YAH. Interclass correlation coefficients were generated through two-way mixed effects, consistency single rater method.

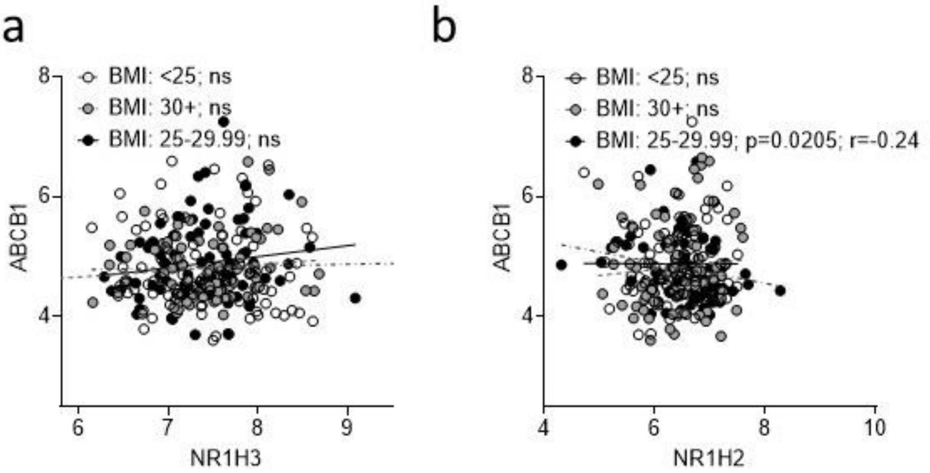


**Supplementary Figure 4: Obesity is not associated with activity of the LXR:Pgp axis in Luminal A and Luminal B patients.** In luminal A and luminal B patients (GSE78958), Pgp is not significantly positively correlated with NR1H3. Data shown are from patients with luminal A (n=209) and luminal B breast cancer (n=43) and grouped according to BMI information (<25 n=85; 25-29.99 n=74; ≥30 n=93) and analysed using Spearman correlation.
